# The N-terminal domain of the human mitochondrial helicase Twinkle has DNA binding activity crucial for supporting processive DNA synthesis by Polymerase γ

**DOI:** 10.1101/2022.11.10.516034

**Authors:** Laura C. Johnson, Anupam Singh, Smita S. Patel

**Affiliations:** Department of Biochemistry and Molecular Biology, Rutgers University, Robert Wood Johnson Medical School, Piscataway, NJ 08854, USA; Graduate School of Biomedical Sciences at the Robert Wood Johnson Medical School of the Rutgers University, USA

**Keywords:** mitochondria, replication, helicase, Twinkle, DNA polymerase, replisome, hexameric helicase, mitochondrial diseases

## Abstract

Twinkle is the ring-shaped replicative helicase within the human mitochondria with high homology to bacteriophage T7 gp4 helicase-primase. Unlike many orthologs of Twinkle, the N-terminal domain (NTD) of human Twinkle has lost its primase activity through evolutionarily acquired mutations. The NTD has demonstrated no observed activity thus far, hence its role has remained unclear. In this study, we have biochemically characterized the isolated NTD and C-terminal domain with linker (CTD) to decipher their contributions to the activities of the full-length (FL) Twinkle. This novel CTD construct hydrolyzes ATP, has weak DNA unwinding activity, and assists Polγ-catalyzed strand-displacement synthesis on short replication forks. However, CTD fails to promote multi-kilobase length product formation by Polγ in rolling-circle DNA synthesis. Thus, CTD retains all the motor functions but struggles to implement them for processive translocation. We show that NTD has DNA binding activity, and its presence stabilizes Twinkle oligomerization. The CTD oligomerizes on its own, but loss of NTD results in heterogeneously-sized oligomeric species. The CTD also exhibits weaker and salt-sensitive DNA binding compared to FL Twinkle. Based on these results, we propose that NTD directly contributes to DNA binding and holds the DNA in place behind the central channel of the CTD like a ‘doorstop’, preventing helicase slippages and sustaining processive unwinding. Consistent with this model, mtSSB compensate for the NTD loss and partially restore kilobase length DNA synthesis by CTD and Polγ. The implications of our studies are foundational for understanding the mechanisms of disease-causing Twinkle mutants that lie in the NTD.

## Introduction

Twinkle is the replicative helicase of the human mitochondria, discovered through its sequence homology to bacteriophage T7 gp4 helicase-primase in linkage studies of mitochondrial-related disease, autosomal dominant progressive external ophthalmoplegia (adPEO) (1). Many point mutations of Twinkle have been identified in association with clinical presentations of a number of varied mitochondrial-related neuromuscular diseases, from the aforementioned adPEO to mitochondrial DNA depletion syndrome, spinocerebellar ataxia and certain presentations of Parkinson’s symptoms (2–5). These mutations disturb mitochondrial DNA replication and generally result in age-related mtDNA deletions or depletion (6). The expression of recombinant human Twinkle in bacteria and insect cells has greatly facilitated biochemical and structural studies of Twinkle and reconstituted the mt replisome complex with the partnering proteins the polymerase Polγ, which is a heterotrimer of PolγA and two PolγB subunits, and the mitochondrial single-stranded binding proteins (mtSSB) (7). Twinkle spontaneously organizes into ring-shaped hexamer and heptamer rings (8,9). A recent cryo-EM study reported a high-resolution structure of a disease-mutant of Twinkle in a closed heptameric/octameric ring arrangement (**Fig. 1**) (10). Biochemical studies indicate that Twinkle is a 5’-3’ helicase with poor unwinding activity on its own (7,9,11). However, in concert with Polγ and mtSSB, Twinkle supports creation of kilo-base-sized products in rolling circle DNA synthesis on a minicircle fork (7). Twinkle also demonstrates DNA annealing activity, the biological role of which remains unknown (9,12).

**Figure 1.**
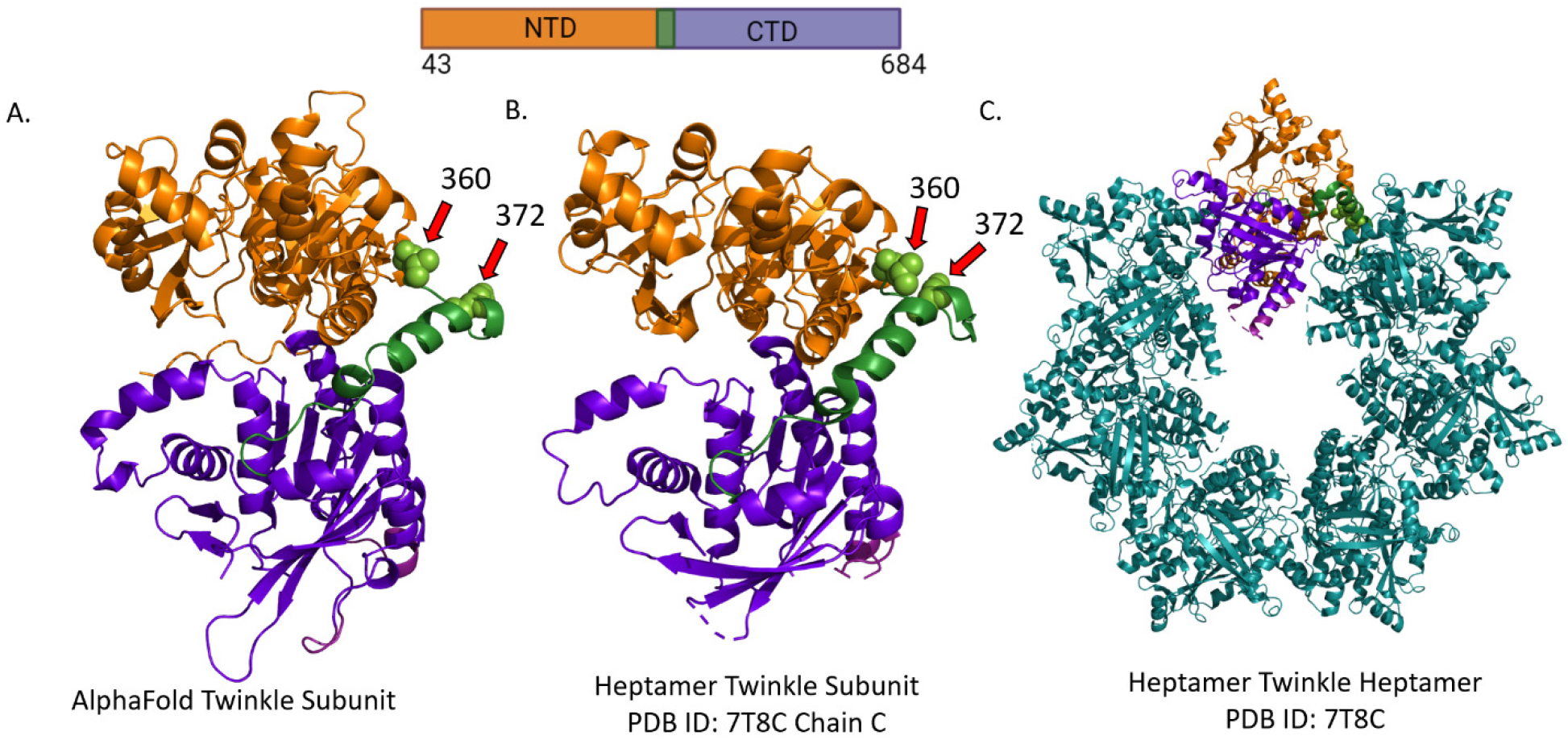
Domain structure of the Twinkle subunit and its organization into a ring **A**. The AlphaFold predicted structure for human Twinkle with the NTD in orange, the CTD in purple and the linker region in green. Deletion position residues 360 and 372 are rendered as spheres and marked with arrows. **B**. The isolated subunit of the heptameric Twinkle (PDB ID: 7T8C). **C**. Cryo-EM structure of the heptameric Twinkle ring (PDB ID 7T8C). The primary structure schematic of the full length construct with included amino acid residues is displayed above with the domains color coded.

Each subunit of Twinkle consists of two roughly equivalently sized globular domain halves divided into an N-terminal domain (NTD) and a C-terminal domain (CTD), separated by a helical linker (**Fig. 1**). The CTD contains the conserved helicase and ATPase motifs (13) and the helix bundle that along with the linker organizes subunit-subunit interactions in ring assembly (10). The NTD includes primase motifs, and many orthologs of the human Twinkle have retained the primase function, but human Twinkle has lost this function due to mutations in the zinc-binding domain and Mg-ion binding site of the primase (13–16). The role of replicative primase in human mitochondria is delegated to the mitochondrial RNA polymerase, POLRMT (17,18). Unlike CTD, the NTD does not participate in inter-subunit interactions but shows many intra-molecular interactions with the CTD (10) (**Fig. S1**). A previous study reported biochemical studies of a partially NTD deleted mutant of Twinkle (314-684 aa), which suggested that NTD is essential for replication functions (19), however, this construct retained much of the NTD proximal to the linker which based on proximity may be involved in intra-molecular contacts (**Fig. S1**) (10). Furthermore, no activity has been demonstrated for the NTD itself, its role in human Twinkle has remained mysterious. The NTD and linker are hotspots for disease-causing mutations (20), suggesting a vital role in Twinkle activity. Deciphering the NTD function is critical for understanding the possible reasons for mutation-related replication defects.

In this study, we have expressed the isolated domains of Twinkle, including the NTD and CTD plus linker, referred to as the CTD here, and carried out a detailed biochemical analysis of these constructs and the FL Twinkle, comparing their oligomerization, DNA binding, ATPase, helicase, and replisome functions. Our studies reveal that NTD has a DNA binding activity, and although CTD can bind DNA on its own, the NTD confers high-affinity binding to FL Twinkle. Additionally, our studies also show that NTD is required to form homogenous oligomers of Twinkle. Both DNA binding and proper oligomerization of Twinkle are essential for processive unwinding to facilitate production of multi-kilo base lengths of DNA through strand-displacement DNA synthesis by Polγ. We propose a door-stop model to explain the role of NTD in processive DNA unwinding.

## Results

### Expression of NTD and CTD protein domains of human Twinkle

To create the isolated domains of Twinkle, we used the alphafold predicted structure of Twinkle, which matches the recently determined structure of Twinkle mutant heptamer (10,21). The experimental and predicted structure of the Twinkle subunit show that the NTD ends around amino acid residue 360, the linker helix region lies between 360-394, and the CTD is formed between 394-684 (**Fig. 1**). Accordingly, we cloned the sequence corresponding to amino acid residues 43-372 to make the NTD construct (35,596 Da) lacking the linker domain; the 1-42 aa is the predicted mitochondrial targeting sequence. This NTD construct was reported in a previously published study of Twinkle (19), while the minimal CTD construct (36,650 Da) that contained amino acids from 360-684, which includes the CTD and the linker region only, is novel. The SUMO-fusion constructs of NTD, CTD, and FL Twinkle were expressed in bacteria and purified as soluble proteins without the SUMO tag. The NTD was expressed abundantly compared to the CTD and FL Twinkle. We have reported the bacterial expression and purification of wild-type FL Twinkle previously (12), but the modified protocol described here produces soluble Twinkle without contaminating nucleic acid, chaperone, or exonuclease activity (**Fig. S2**).

### Size-exclusion gel filtration analysis of CTD, NTD, and full-length Twinkle

The oligomeric states of purified FL Twinkle, NTD, and CTD proteins were assessed using size-exclusion gel-filtration chromatography (**Fig. 2**). Based on the elution volumes of the protein markers (**Fig. 2A**), FL Twinkle elutes close to a heptamer with a small peak corresponding to protein aggregates (**Fig. 2B**). The CTD elution peak was broader and centered around a larger oligomeric species (~12 subunit) (**Fig. 2C**). The NTD eluted as a monomer (**Fig. 2D**), even at the high concentrations of NTD used in this experiment (~80 µM). These results indicate that oligomerization of FL Twinkle is driven mainly by the CTD, though the broader elution peak of the CTD construct relative to FL Twinkle indicates that NTD regulates the oligomeric structure of Twinkle by preventing higher order species and promoting hexamer/heptamer formation.

**Figure 2.**
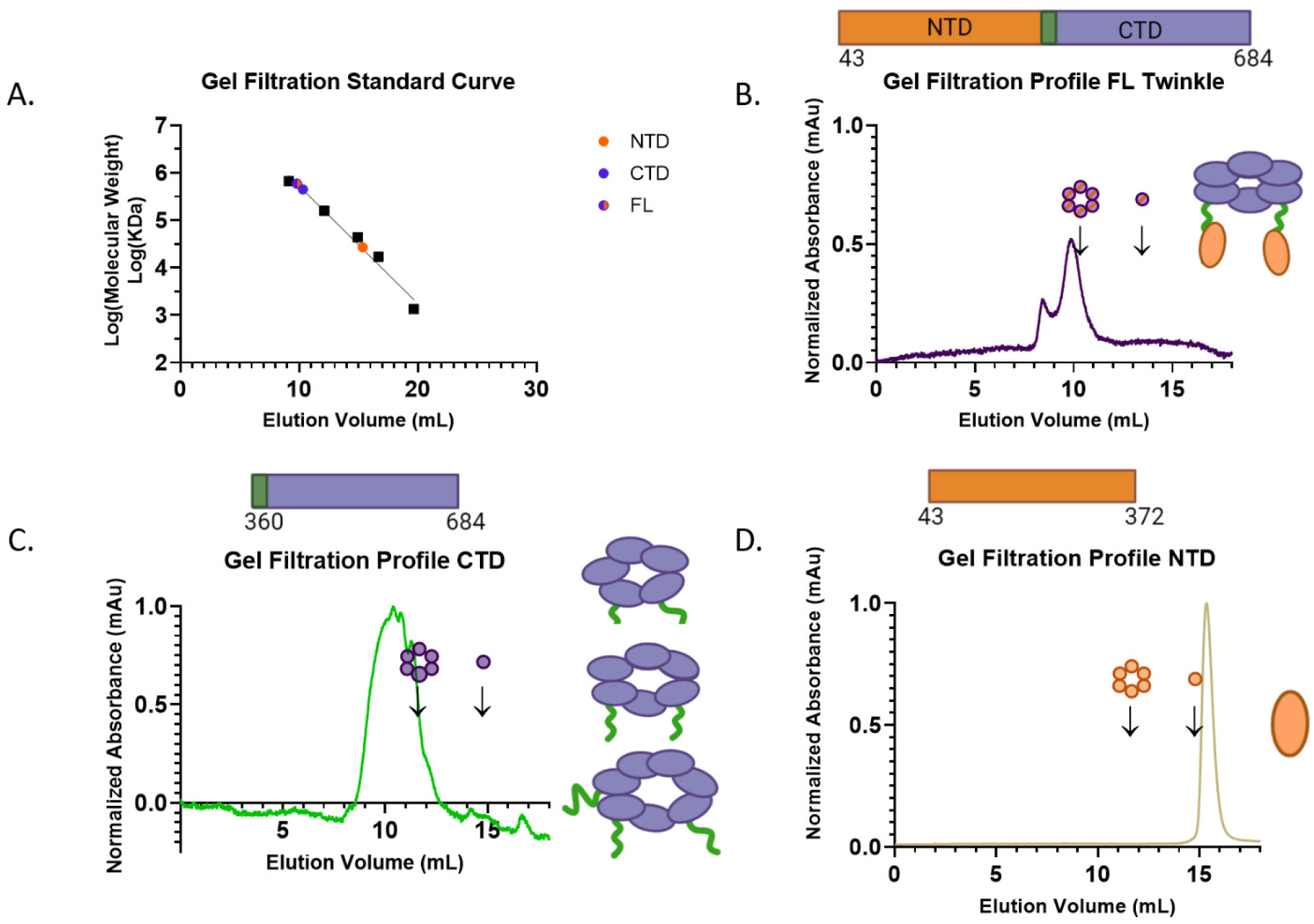
Gel-filtration analysis to assess the oligomerization of purified FL Twinkle, CTD, and NTD constructs. Samples were injected in Superdex 200 Increase 10/300 GL Cytiva column with respective running at 0.3 mL/min and peak elution volume monitored via 280 nm absorbance. **A**. Calibration curve shows the elution volumes of the Biorad gel filtration protein standards as black squares, the elution volume of each Twinkle construct is displayed as appropriately colored circles **B**. Full length human Twinkle elution profile monitored using absorbance at 280 nm. Expected locations of hexameric and monomeric Twinkle are marked as determined by the calibration curve. **C**. Twinkle CTD elution profile with the expected locations of hexameric and monomeric CTD. **D**. Twinkle NTD elution profile with the expected locations of hexameric and monomeric NTD. The oligomerization state(s) interpreted as mostly likely present in each spectrum based on the elution profile is displayed to the right, while the primary structure schematic of each construct with included amino acid residues is displayed above each respective graph.

### FL Twinkle and CTD bind single-stranded DNA in a length-dependent manner

Ring-shaped helicases generally bind single-stranded (ss) DNA in the central channel, and the structure of the archetypal T7 gp4 helicase shows that each subunit contacts two nucleotides; thus, the minimal DNA binding length is 12 nt per hexamer (22,23). We made a series of ssDNA ligands of random sequence from 8-nt to 30-nt (**Table S1**) to determine the DNA binding affinity and the minimal DNA length required to bind FL Twinkle and CTD. Each DNA contained a fluorescein moiety at the 3’-end, which enabled us to measure the K_D_ values using fluorescence anisotropy titrations, which unlike mobility shift assays are not disruptive and measures binding under equilibrium conditions. An increasing protein concentration was added to a constant amount of fluorescein-labeled ssDNA, and the resulting increase in anisotropy were plotted as binding curves that were fit to a 1:1 binding equation to obtain the K_D_ values (**Fig. 3**). The K_D_ values were measured in 50 mM Tris-acetate buffer under two salt conditions, one without added NaCl and one with 50 mM NaCl (**Table 1**).

**Table 1.** Equilibrium dissociation constants (K_D_) for FL Twinkle, CTD, and NTD constructs binding to single-stranded DNAs of increasing lengths determined from fluorescence anisotropy based titrations fit to a one site binding hyperbolic equation. The * represents the tight binding phase K_D_ value from fit to sum of two hyperbolas.

| | $K_D$ (nM) no salt added | | |
| --- | --- | --- | --- |
| DNA length | FL Twinkle | CTD | NTD |
| 8 | $9.0 \pm 4.9$ | $116.1 \pm 41.1$ | - |
| 12 | $5.2 \pm 1.0$ | $21.7 \pm 3.0$ | $140.8 \pm 25.2$ |
| 14 | $3.3 \pm 0.7$ | $8.8 \pm 1.5$ | $120.6 \pm 25.8$ |
| 16 | $2.8 \pm 0.7$ | $5.2 \pm 1.4$ | $104.5 \pm 23.8$ |
| 20 | $1.9 \pm 0.$ | $2.2 \pm 0.7$ | $18.0 \pm 7.0^*$,<br>$1204 \pm 643$ |
| 30 | $1.6 \pm 0.3$ | $1.7 \pm 0.5$ | $17.5 \pm 4.0^*$,<br>$734 \pm 449$ |

**Table 1** Equilibrium dissociation constants ( $K_D$ ) for FL Twinkle, CTD, and NTD constructs binding to single-stranded DNAs of increasing lengths determined from fluorescence anisotropy based titrations fit to a one site binding hyperbolic equation.
|  | K<sub>D</sub> (nM) + 50 mM NaCl |  |  |
| --- | --- | --- | --- |
| DNA length | FL Twinkle | CTD | NTD |
| 8 | 9.8 ± 11.8 | - | - |
| 12 | 53.1 ± 17.4 | - | - |
| 14 | 39.0 ± 16.2 | 128.1 ± 83.5 | - |
| 16 | 22.6 ± 5.1 | 116.5 ± 44.7 | - |
| 20 | 8.8 ± 1.3 | 50.0 ± 11.0 | 444.3 ± 258.9 |
| 30 | 2.6 ± 0.5 | 4.5 ± 0.9 | 264.1 ± 61.6 |

**Figure 3.**
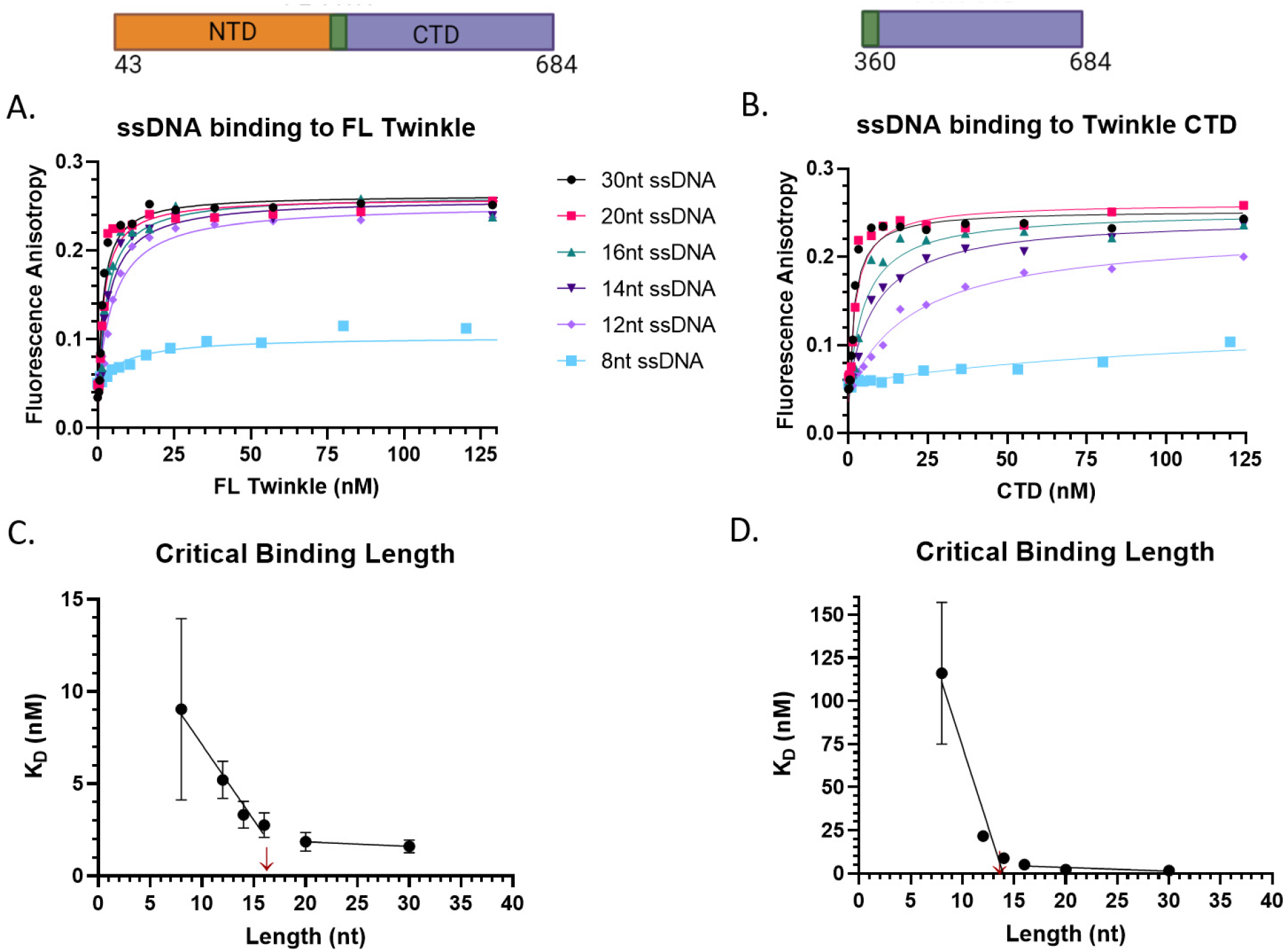
Length dependent binding of single-stranded DNAs to FL Twinkle and CTD. The fluorescence anisotropy-based titrations were carried out in buffer containing 50 mM Tris acetate pH 7.5, 10 % glycerol, 0.05 % Tween 20, 0.5 mM DTT using 2.5 nM ssDNA (8 nt, 12 nt, 14 nt, 16 nt, 20 nt, 30 nt) titrated with hexameric concentrations of FL Twinkle and CTD. **A**. Length dependence binding titration curves for FL Twinkle on a series of ssDNA lengths **B**. Length dependence binding titration curves for CTD on a series of ssDNAs. The equilibrium dissociation constant (K_D_) versus ssDNA length for FL Twinkle **C**. and CTD **D**. The primary structure schematic of each construct with included amino acid residues is displayed above each respective binding titration graph. Each curve fit to one site binding hyperbolic equation, and standard error of fit are shown.

FL Twinkle and CTD bind DNA of various lengths with nM K_D_ values, but overall, the affinity of CTD for the DNA is weaker than FL Twinkle, particularly DNAs from 8-nt to 16-nt in length (**Table 1**). Each protein shows a length dependency. The 8-nt DNA binds weakly to both FL Twinkle and CTD but the affinities increase with increasing ssDNA length (**Fig. 3A,B**). A plot of K_D_ versus DNA length indicate that the critical DNA binding length is 14-16 nt (**Fig. 3C,D**). If each Twinkle subunit binds to 2-nt of DNA as with T7 gp4 helicase, then this result suggests that both CTD and FL Twinkle assume a hexameric/heptameric structure in the presence of DNA without additional cofactors.

### DNA binding to CTD is more salt sensitive than FL Twinkle

Adding 50 mM NaCl weakened the DNA binding affinities of both CTD and FL Twinkle (**Fig. 4**). However, salt addition had a more pronounced effect on CTD than the FL Twinkle. For example, under no NaCl conditions, CTD binds to 12-nt with K_D_ of 22 nM, which is 4-fold weaker than FL Twinkle K_D_ (~ 5 nM). In 50 mM NaCl, CTD showed barely any binding to 12-nt DNA whereas FL Twinkle bound with 50 nM K_D_. Salt addition also reduced the binding affinity of the longer 14-20 nt oligos by 14-22-fold for CTD as opposed to 5-10 for FL Twinkle. These DNA binding studies demonstrate that CTD can bind DNA on its own, but NTD is necessary for stable binding. Thus, NTD directly or indirectly contributes to DNA binding in FL Twinkle.

**Figure 4.**
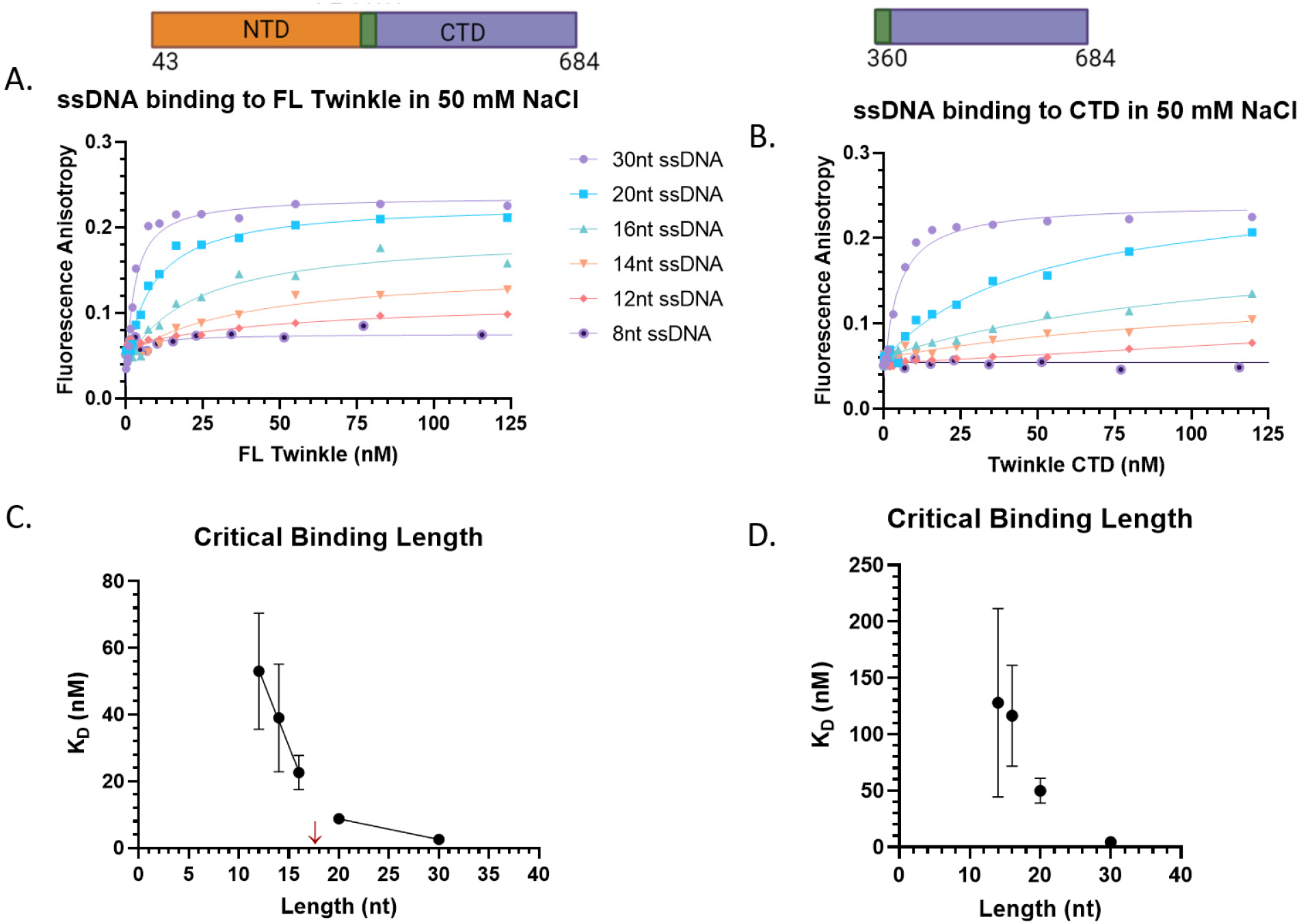
Single-stranded DNA binding to FL Twinkle and CTD in 50 mM NaCl. The fluorescence anisotropy-based titrations were carried out in same buffer as in Figure 3 with 50 mM NaCl added. Each curve was fit to one site binding hyperbolic equation. **A**. Length dependence binding titration curves for FL Twinkle for the series of ssDNA lengths **B**. Length dependence binding titration curves for CTD for the series of ssDNAs. The equilibrium dissociation constant (K_D_) versus ssDNA length for FL Twinkle **C**. and CTD **D**. The primary structure schematic of each construct with included amino acid residues is displayed above each respective binding titration graph. Each curve fit to one site binding hyperbolic equation, and standard error of fit are shown.

### Twinkle NTD binds to ssDNA

To determine whether NTD contributes directly to DNA binding, we used fluorescence-anisotropy based titrations to measure the K_D_ values of NTD for the same series of 8-nt to 30-nt length DNAs we used to evaluate FL Twinkle and the CTD. A previous study had used gel mobility shift assay and concluded that NTD does not bind to DNA (19). Interestingly, in our hands, NTD has a DNA binding activity that can be quantified by equilibrium titrations (**Fig. 5**). The NTD displayed negligible binding to the 8-nt DNA, but the 12-nt to 30-nt DNAs showed nM K_D_ values with affinities increasing with increase in DNA length. Interestingly, the 20-nt and 30-nt DNAs showed a biphasic behavior in binding titrations (**Fig. S3**). The high-affinity binding mode displays K_D_ values around 20 nM and the low-affinity mode around 730 to 1200 nM (**Table 1**). The biphasic behavior suggests that two NTDs are involved in binding 20-30 nt DNA.

**Figure 5.**
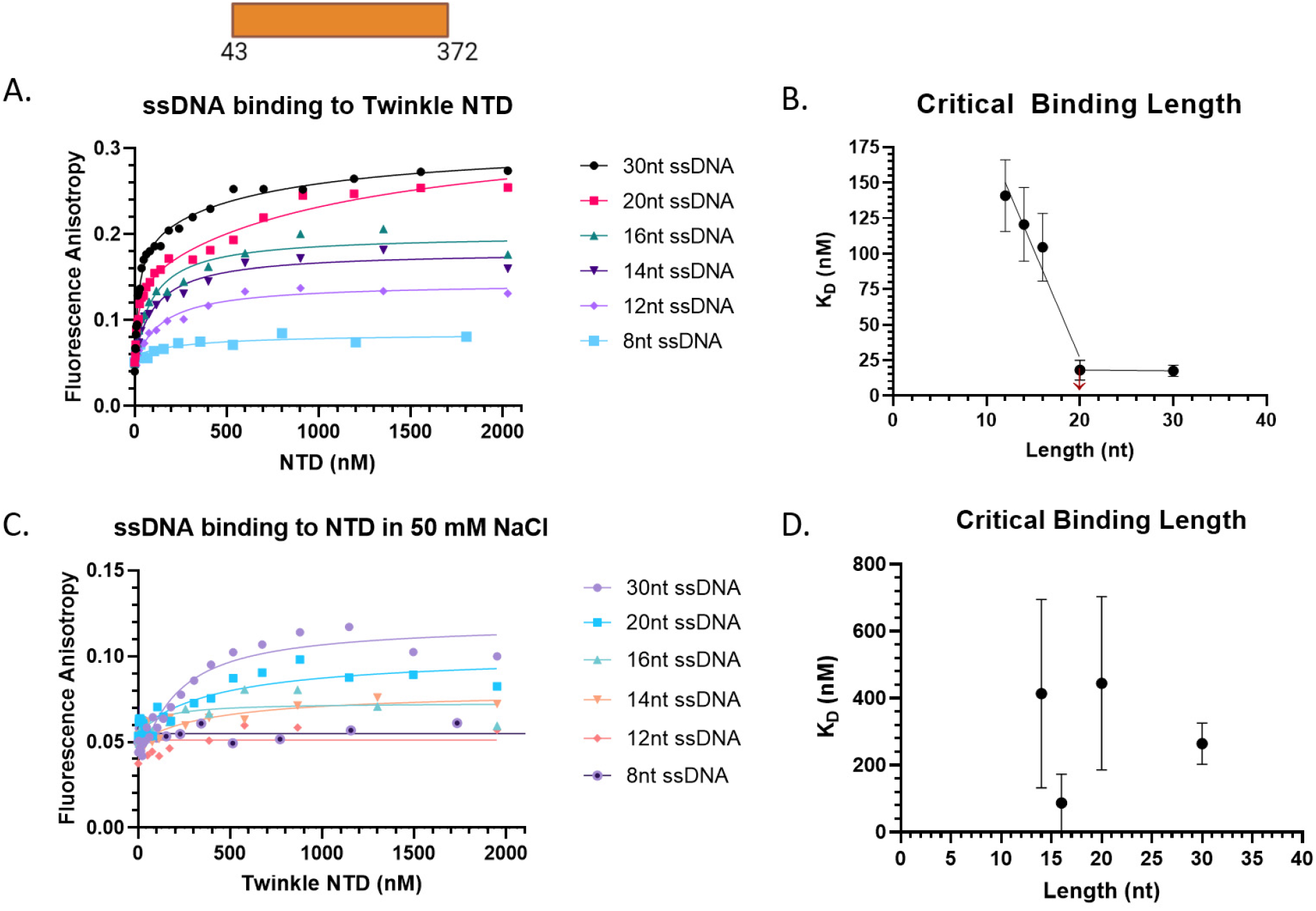
Length dependent binding of single-stranded DNAs to Twinkle NTD. The fluorescence anisotropy-based titrations were carried out as detailed in figure 3 and figure 4 legends. **A**. DNA binding titration curves of NTD for a series of ssDNA in no salt added conditions. **B**. NTD K_D_ versus ssDNA length in no salt added conditions. **C**. DNA binding curves for NTD in 50 mM NaCl conditions. **D**. NTD K_D_ versus ssDNA length in 50 mM NaCl conditions. Each curve fit to one site binding hyperbolic equation except for 30-nt and 20-nt ssDNA in no salt added conditions, which were fit to two site binding hyperbolic equation, and standard error of fit are shown.

The DNA binding activity of NTD was acutely salt sensitive. The 50 mM NaCl competed effectively with the binding of short DNAs from 8-16 nt, and the salt decreased the affinity of the 30-nt DNA by 15 fold, from ~20 nM to ~260 nM. These results demonstrate that NTD has a DNA binding activity that can contribute directly to the overall high affinity and stability of the FL Twinkle on DNA.

### The CTD hydrolyzes ATP and unwinds duplex DNA, but less efficiently than FL Twinkle

The ATPase motifs are present in the CTD, and these motifs form an active site at the subunit interface in the ring structure (10). The ATPase activity is essential for supporting translocation and unwinding activities of the helicase. The time course of ATP hydrolysis shows that both FL Twinkle and CTD hydrolyze ATP in the absence and presence of DNA (**Fig. 6A**). Compared to FL Twinkle, the ATP hydrolysis rate of CTD was only two-fold lower (**Fig. 6B**). Thus, NTD loss lowers the ATPase activity of the CTD, but the impairment is moderate. Adding M13 ssDNA increases the ATPase rate of both FL Twinkle and CTD by two-fold. These results indicate that the isolated CTD can assemble into ATPase-active oligomers both in the presence and absence of DNA.

**Figure 6.**
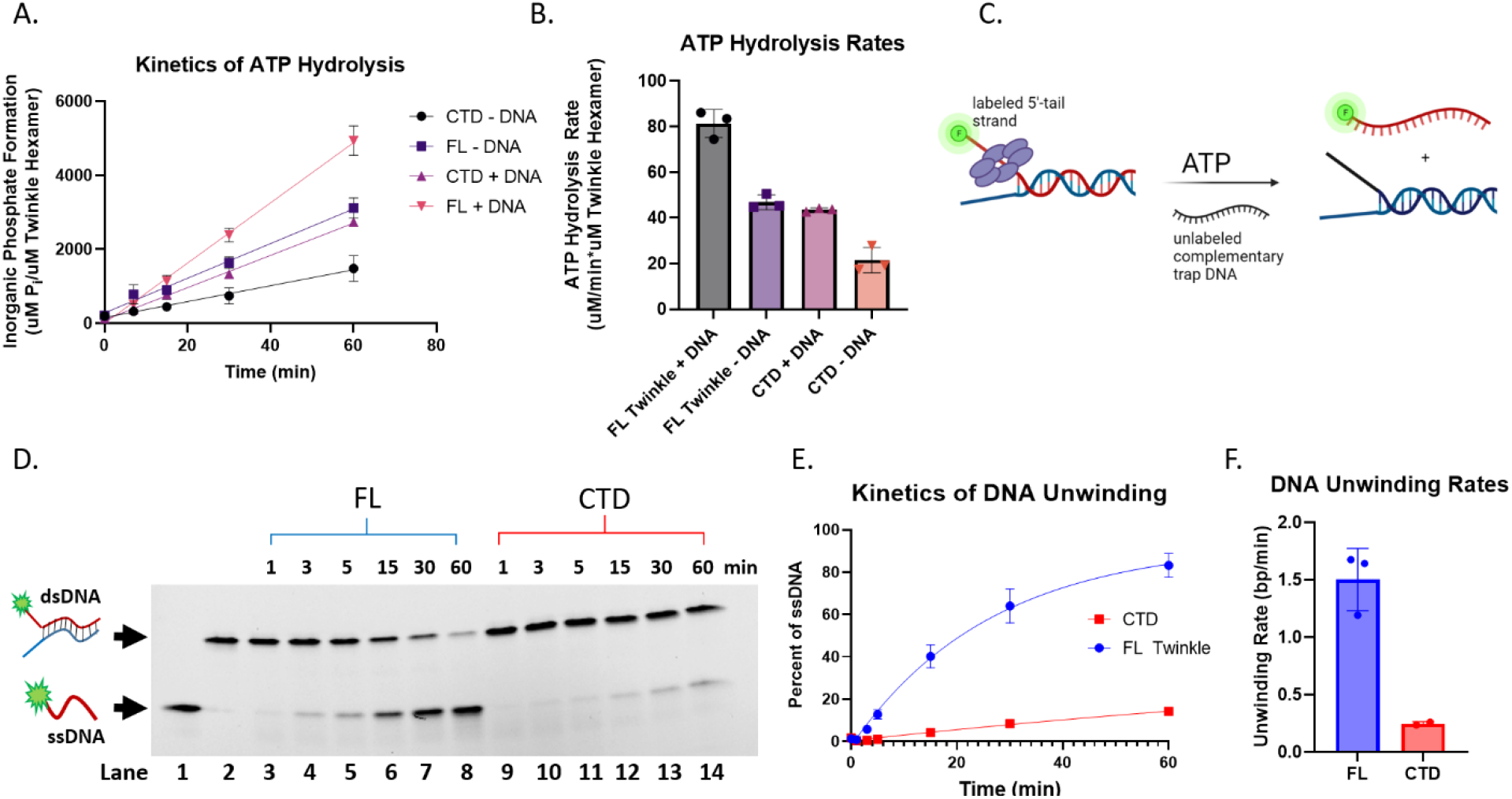
ATP hydrolysis and DNA unwinding kinetics of CTD and FL Twinkle. **A**. Time courses of ATP hydrolysis reaction for CTD and FL Twinkle with and without DNA. ATP hydrolysis was measured using 30 nM CTD or FL Twinkle (hexamer) with and without 2.5 nM M13 ssDNA molecules in 8 mM magnesium acetate and 1 mM ATP spiked with [γ-_32_ P] ATP. **B**. The ATP hydrolysis rates from three replicate kinetics from panel **A**. are shown with the standard deviations. **C**. Schematics of the unwinding reaction showing 5’-ssDNA tail bound Twinkle catalyzing the release of fluorescent 5’-tail DNA in the presence of ATP and trap DNA. The unwinding assay was carried out using same buffer with 10 nM forked DNA, 4.5 mM ATP, 8 mM magnesium acetate, 55.5 nM Twinkle hexamer, 280 nM trap DNA. **D**. Representative image of the 4-20 % native polyacrylamide gel showing the time course of ssDNA formation by FL Twinkle and CTD. Lane 1: free 5’-tail ssDNA, lane 2: free forked DNA, and lanes 3-8, and 9-14 show time courses of forked DNA unwinding by FL Twinkle and CTD, respectively. **E**. DNA unwinding kinetics showing proportion of 5’-tail release for the FL Twinkle and CTD fit to an exponential equation.**F**. The rate of DNA unwinding graphed with individual replicates as points and error bars.

Twinkle is a 5’-3’ helicase and assembles on the 5’-ssDNA tail to initiate DNA unwinding of the duplex region (9,11,24). We tested the helicase activity of FL Twinkle and CTD using a 40-bp forked DNA (**Table S2**) under single-turnover conditions (**Fig. 6C**). The 40-bp fork DNA contained a 28-nt 3’-tail and a 35-nt 5’-tail labeled with fluorescein to monitor DNA strand separation using a native gel assay. The helicase reactions were carried out by incubating Twinkle with the forked DNA and adding ATP, Mg^2+^, and an excess of trap DNA to capture the unwound strand (unlabeled lower strand) and the free/dissociated Twinkle. FL Twinkle unwinds the fork DNA at a rate of 1.5 bp/min, but the CTD shows a weak DNA unwinding activity with a rate of 0.25 bp/min (**Fig. 6D-F**). This indicates that deletion of NTD has a more drastic effect on the helicase activity than the ATPase activity. Thus, the isolated CTD can assemble into ATPase active oligomers in the absence of NTD, but NTD is necessary to functionally couple the ATPase activity to DNA unwinding.

### Twinkle CTD supports strand-displacement DNA synthesis by Polγ on a short replication fork DNA

Twinkle, Polγ, and mtSSB have been shown to catalyze rolling circle synthesis (7). Herein, we used a short 40-bp replication fork consisting of a 30-nt primer to measure wild-type Polγ’s strand-displacement DNA synthesis activity with and without Twinkle (**Table S2, Fig. 7A**). We used wild-type Polγ, which has proofreading 3’-5’ exonuclease activity, because exo-Polγ has been shown to have intrinsic strand displacement synthesis activity in the absence of Twinkle (25,26). The 45-nt 5’-tail of the forked DNA is long enough to bind one Twinkle oligomer; hence, mtSSB was not added to the reactions. In the absence of Twinkle, Polγ fills the 4-nt ssDNA gap between the 3’-end of the primer and the start of the 40-bp duplex region in the replication fork but does not carry out strand-displacement DNA synthesis (**Fig 7B, lane 1**). When FL Twinkle was added, Polγ could strand-displace and fully extend the primer to the full-length product (**Fig. 7B, lanes 6-9**), demonstrating that Twinkle and Polγ can catalyze DNA unwinding-synthesis without mtSSB on a short fork DNA. We also observed an accompanying excision reaction, which degraded part of the primer at the start. Interestingly, despite its previously characterized poor unwinding activity (**Fig.6D-F**), the CTD could catalyze unwinding-synthesis with Polγ on the short replication fork DNA. The primer extension yield in the CTD reactions was 3-fold lower compared to FL Twinkle (**Fig. 7C**), and the DNA synthesis rate of Polγ and CTD was 2-fold lower (**Fig. 7D**). These results indicate that CTD can functionally couple with Polγ to catalyze strand displacement synthesis, albeit with lesser efficiency.

**Figure 7.**
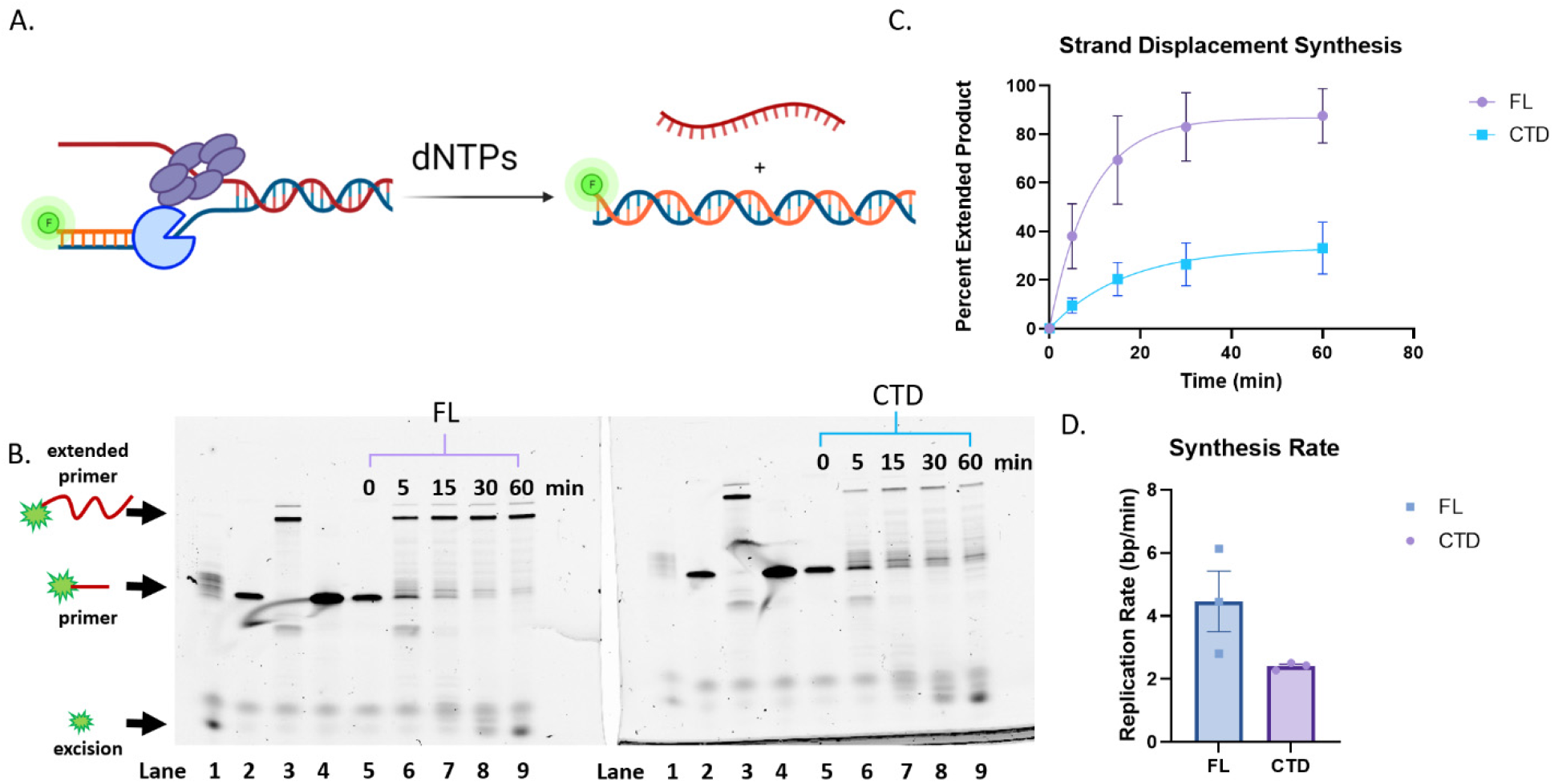
Strand displacement DNA synthesis activity of Polγ with and without FL Twinkle or CTD. **A**. Schematic shows the strand displacement DNA synthesis assay with fluorescein labeled primer to monitor primer extension by the denaturing gel assay. Reactions were carried out with 300 nM Twinkle hexamer or CTD hexamer, 100 nM forked DNA, 200 nM Polγ, 300 µM dCTP, 100 µM remaining dNTPs, and 4 mM ATP. **B**. Representative image of the 15 % TBE-Urea gel resolving the starting primer and extended primer from DNA synthesis. Samples from reactions with FL Twinkle or CTD were loaded in identical lanes in the two gels. Lane 1: fork DNA + Polγ reaction, lane 2: forked DNA alone, lane 3: Polγ + primer-template reaction, lane 4: free primer, lanes 5-9: reactions with Polγ and FL Twinkle or CTD. **C**. Proportion of the extended primer with Polγ and FL Twinkle or CTD (error bars three replicates). The kinetics fit to a single exponential equation to provide a composite rate of DNA synthesis over the 40-bp duplex (solid lines). **D**. Rates of DNA synthesis in bp/min for FL Twinkle and CTD with individual replicates as points and error bars representing standard deviation.

### Twinkle CTD does not support rolling circle DNA synthesis by Polγ

The entire mitochondrial genome of ~16 kb is copied by the leading strand mt replisome complex consisting of Twinkle, Polγ, and mtSSB (27). Such long stretch DNA synthesis can be assessed *in vitro* using a rolling circle DNA synthesis on a 70-bp minicircle DNA (7) which we used to assess the replication activity of Polγ with CTD (**Fig. 8A**). The radioactively labeled reaction products were resolved on an alkaline agarose gel to assess the size and yields of the DNA products (**Fig. 8B**)(**Table S3**). As shown by the markers, the alkaline agarose gel separates DNA products from a few hundred bases in length to > 10 kilo-bases. Polγ alone does not catalyze rolling circle synthesis on its own and no detectable DNA products were observed in those reactions. However, in the presence of FL Twinkle, Polγ was able to make >10 kb sized products with an average size around 4 kb (**Fig. 8C**). This result demonstrates that FL Twinkle and Polγ can cooperatively catalyze processive strand displacement DNA synthesis even in the absence of mtSSB. Substituting FL Twinkle with CTD reduced DNA synthesis drastically (**Fig. 8D**); faint bands of 0.5 to 1 kb length products were detected but in meager yields (**Fig. 8D,F**). These results indicate that CTD requires the activity of the NTD to catalyze strand-displacement DNA synthesis over long DNA stretches.

**Figure 8.**
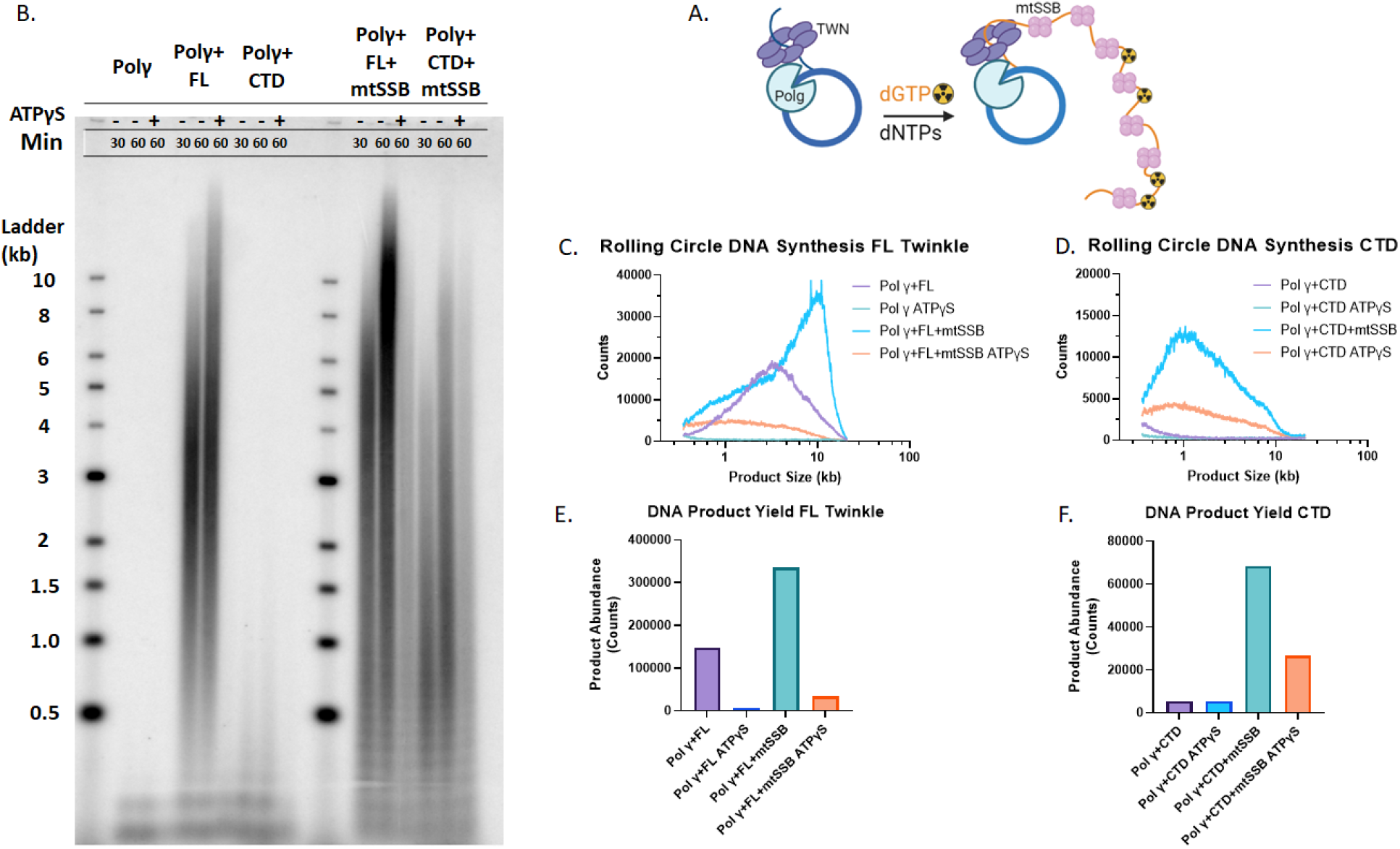
Rolling circle DNA synthesis on the 70-bp minicircle fork DNA. **A**. Schematic shows rolling circle DNA synthesis on the 70-bp minicircle forked DNA with Twinkle, Polγ, and mtSSB. Reactions were carried out using 20 nM FL Twinkle or CTD hexamer, 20 nM wild-type Polγ, 250 nM mtSSB when added, 10 nM 70 bp minicircle forked DNA, 2 mM ATP, 250 μM dNTPs, 25 μM dGTP spiked with [α-^32^P]dGTP at 37°C. **B**. Representative image of an 0.8 % alkaline agarose gel showing the DNA products from the rolling circle DNA synthesis with the DNA size marker ladder. **C**. Quantified counts of the 60 min reaction as a function of product length with FL Twinkle and Polγ with and without mtSSB and ATPγS. Product lengths were determined from the calibration curve generated from the DNA ladder run in the same gel. **D**. Quantified counts of the 60 min reaction as a function of product length with CTD and Polγ with and without mtSSB and ATPγS. **E**. Total DNA product counts for each reaction containing FL Twinkle and Polγ with and without mtSSB and ATPγS **F**. The total product counts for each reaction containing CTD and Polγ with and without mtSSB and ATPγS.

The leading strand DNA synthesis reaction by the mt replisome produces long stretches of ssDNA covered with mtSSB *in vivo* (28). The *in vitro* rolling circle DNA synthesis also produces long stretches of ssDNA (1 kb to >10 kb) that must be covered with mtSSB to prevent secondary structure formation and off target Twinkle or Polγ binding. Thus, addition of mtSSB increased the average sizes of DNA synthesized by FL Twinkle and Polγ from 4 kb to 11 kb and increased the yield of DNA synthesis by 2-fold (**Fig. 8C,E**)(**Table S3**). Remarkably, adding mtSSB stimulated the CTD and Polγ reactions by a significant 12-fold, and increased the observed size of synthesized DNA from a few hundred bases to ~10 kb lengths with an average size of ~ 2 kb (**Fig. 8D,F**)(**Table S3**). This dramatic increase in DNA synthesis processivity with CTD and mtSSB was unexpected. To tease out the contribution of CTD versus mtSSB to DNA unwinding, we substituted ATP with ATPγS, which should inhibit the contribution of the motor activity from the CTD but not the energetically passive activity of the mtSSB (**Fig. 8B,D**). These experiments show that about one-third of synthesis products are from Polγ and mtSSB catalysis, and the remaining two-thirds are from CTD, Polγ, and mtSSB catalysis (**Fig. 8F**) (**Table S3**). Thus, mtSSB stimulates the unwinding activity of CTD and partially compensates for the loss of NTD by binding to the nascent DNA behind the CTD.

## Discussion

In this study, we have determined the role of the N-terminal domain of Twinkle by biochemically characterizing the isolated NTD and CTD domains and comparing their activities to FL Twinkle. Human Twinkle is homologous to bacteriophage T7 gp4 helicase-primase, whose NTD harbors a primase function necessary for Okazaki fragment synthesis. The prevailing model of human mitochondrial DNA replication is the asynchronous strand displacement mechanism, where the heavy strand of the mtDNA is copied continuously from RNA primers made by the mitochondrial RNA polymerase POLRMT without Okazaki fragment synthesis (18,27–30), although alternative models exist (31). Because lagging strand synthesis does not occur concomitant with leading strand synthesis, frequent priming is unnecessary, which could be the reason for the evolutionary loss of primase function in human Twinkle. This raises the question of why the NTD, constituting nearly half of the protein by mass and volume, is maintained in human Twinkle. A previous study showed that NTD is necessary, and its partial deletion impairs rolling circle DNA synthesis with Polγ and mtSSB (19). Because no activity was demonstrated for the NTD, its precise role was unclear.

In this study, we show that Twinkle NTD has a DNA binding activity, and it also lends stability and uniformity to Twinkle oligomers. We show that while the CTD and linker region construct can oligomerize, it elutes as a broad peak with higher order species from gel-filtration compared to FL Twinkle which elutes as a hexamer/heptamer. The recent structure of a disease-causing Twinkle mutant in heptameric/octameric form and the alphafold subunit structure of Twinkle shows that NTD makes many interactions with the linker region (**Fig. S1**), which supports NTD’s role in supporting the ring structure. The linker and NTD interface are also where many disease-causing mutations are found (**Fig. S1**). Many of these mutants disrupt hexamer formation and generate larger oligomeric rings or broken rings (10,32,33).

The defect in DNA binding due to NTD deletion has a more significant effect on the helicase function of CTD rather than its ATPase activity. Because the ATPase site lies at the subunit interface, the formation of oligomer would be sufficient for ATP hydrolysis. On the other hand, the helicase function requires coordination between DNA binding-release steps and ATP binding-hydrolysis steps in each subunit (34,35). Hence, any defect in DNA binding activity can decouple the ATPase activity from the motor function resulting in poor helicase function. For example, T7 gp4 binds DNA more weakly in the presence of ATP compared to dTTP; and although, ATP is hydrolyzed by T7 gp4, it does not support DNA unwinding, (36,37). Processive unwinding with little slippage was observed with dTTP in single molecule unwinding experiments with T7 gp4, whereas repeated unwinding and slippage events were observed with ATP (38). Thus, in Twinkle, the loss in DNA binding energy due to lack of NTD could increase helicase slippage events explaining the more significant impairment in unwinding function compared to the ATPase activity. Interestingly, CTD and Polγ together were able to catalyze DNA unwinding-synthesis reaction on the short fork with product yields higher than the unwinding reaction with CTD alone. Thus, concomitant DNA synthesis by Polγ can partially restore the unwinding defect caused by NTD deletion. Such cooperativity has been observed in T7 replisome, where the DNA polymerase accelerates the unwinding activity of T7 gp4 (34,35,39).

Although CTD and Polγ could catalyze strand-displacement synthesis on the short replication fork, the two could not catalyze kilo-base length DNA products in rolling circle DNA synthesis as with FL Twinkle. This demonstrates the critical need for the NTD in processive unwinding to support genome-sized DNA products in leading strand synthesis. We were intrigued when mtSSB was able to partially restore the synthesis of long DNA products by Polγ and CTD. The mtSSB also stimulated strand-displacement synthesis activity of the FL Twinkle but more significantly activated the CTD reactions. The tetrameric mtSSB binds ~ 30 nt ssDNA with a high affinity (40) and is expected to coat the nascent ssDNA emerging from the central channel of the Twinkle molecule unwinding DNA at the fork junction (28). The CTD subunits are undergoing significant conformational changes unwinding the base pairs at the fork junction and moving the DNA in the central channel. The mtSSB binding behind Twinkle could minimize the backward slippages of the CTD and prevent DNA from dissociating (**Fig. 9**). We propose that the DNA binding activity of the NTD has such a role in activating and increasing the processivity of DNA unwinding by Twinkle. The NTD serves as a ‘door-stop’ and holds the DNA emerging from the CTD near the central channel preventing the DNA from dissociating and CTD from slipping backward. When linked to CTD as a part of the FL Twinkle, two or three NTDs can cooperate and bind DNA strongly, although non-specific binding may be necessary so NTDs do not inhibit DNA translocation. Thus, our studies reveal that even though the NTD of the human Twinkle has lost its primase function, it has retained its DNA binding activity, enabling the helicase functions of the CTD.

**Figure 9.**
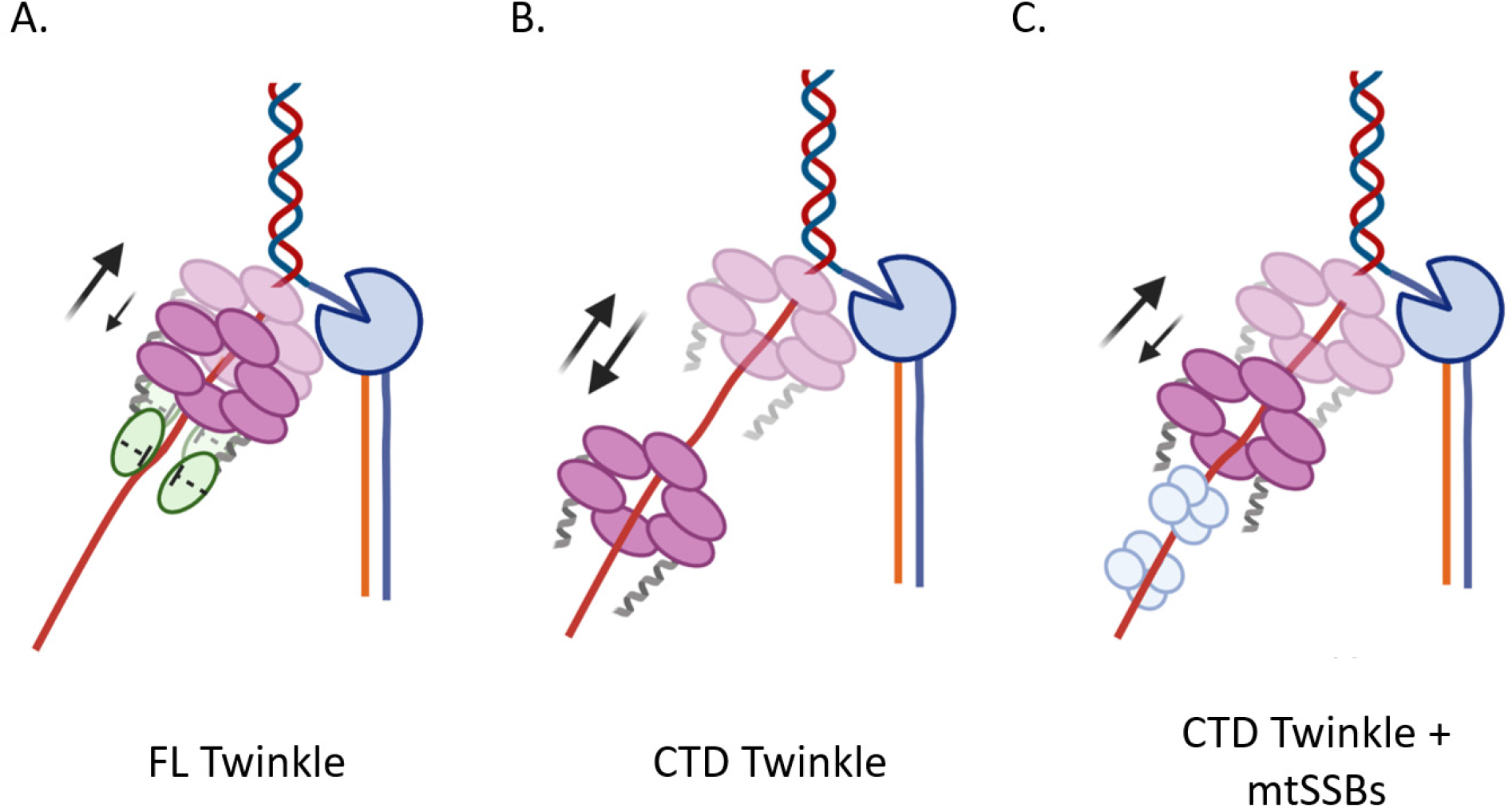
Proposed doorstop mechanism for NTD’s role in supporting processive translocation for DNA unwinding as part of a replisome with Polγ and mtSSBs **A**. The NTD binding to the DNA emerging from the central channel of the CTD domains in FL Twinkle minimize backward motion (down arrow) promoting forward motion (up arrow). **B**. The CTD at the replication fork has more significant backward motion. **C**. The mtSSB molecules binding the DNA behind the CTD ring prevent excessive backward movements

### Experimental procedures

#### Nucleic Acid Substrates

Oligodeoxynucleotides were custom-synthesized with either 5’ or 3’-end fluorescein inverted T, and Biotin modifications and purified by HPLC (Integrated DNA Technologies, Coralville, IA, USA). The 70-nt minicircle DNA was prepared by ligation reaction and assembled into a fork DNA, as described previously (41). DNA concentration was determined from the absorbance at 260 nm and the corresponding molar extinction coefficients.

#### Protein purification

##### PolγA

The catalytic and accessary subunits of human mitochondrial DNA polymerase gamma (PolγA and PolgγB, respectively) were purified as reported previously (42). Histidine tagged PolγA subunit lacking 1-29 putative mitochondrial localization signal sequence and 10 of the 13 sequential Glutamine residues (amino acid residues 43-52) was expressed in Sf9 insect cells. Briefly, Sf9 cells expressing PolγA were lysed by gently stirring the cell pellets in lysis buffer consisting of 70 % of lysis buffer P (0.32 M Sucrose, 10 mM HEPES, pH 7.5, 0.5 % (v/v) NP-40, 3 mM CaCl_2_, 2 mM magnesium acetate, EDTA free protease inhibitor tablet (Roche) and 5 mM DTT). The centrifuged cell lysate was purified using Ni affinity chromatography (HisTrap HP, Cytiva, Marlborough, MA), followed by purification on Heparin column (HiTrap Heparin HP affinity column, Cytiva, MA). The resulting protein was further purified to homogeneity by passing it through Superose 6 gel filtration column (Cytiva, MA).

##### PolγB

Histidine tagged PolγB was expressed in *E*.*coli* Rosetta (DE3) cells. Cell pellets were lysed in 20 mM HEPES pH 8.0, 300 mM KCl, 5 % Glycerol, 0.1 % Triton X-100, 1 tablet of EDTA free protease inhibitor per 50 ml buffer (Roche), 0.1 mg/ml Lysozyme and 10 mM beta mercaptoethanol (βME) using sonication, treated with 0.1 % final concentration of Polyethyleneimine (PEI), and supernatant was applied to a prepacked Ni affinity column (HisTrap HP, Cytiva, MA) pre-equilibrated with Ni buffer (20 mM HEPES pH 8.0, 300 mM KCl, 5 % Glycerol). The bound protein was eluted with a continuous 20 mM-500 mM imidazole gradient. The eluted fractions enriched in PolγB were pooled, diluted to adjust KCl concentration to 60 mM and loaded on a cation exchange chromatography column (HiTrap SP HP Column, Cytiva, MA) pre-equilibrated with SP column low salt buffer containing 20 mM HEPES (pH 7.5), 60 mM KCl, 5 % Glycerol and 5 mM βME. The column was washed with the low KCl buffer to remove loosely bound contaminants. The bound PolγB was then eluted with a continuous gradient of 60 mM to 1 M KCl. The collected fractions with eluted PolγB were analyzed using SDS-PAGE and the fractions with purified protein were pooled. The purified protein was diluted with SP column low salt buffer to achieve final KCl concentration of 100 mM KCl. Purified proteins were concentrated using Amicon® Ultra Centrifugal Filters with MWCO 10,000 (MilliporeSigma, MA). Concentrated proteins were stored at-80°C. To make PolγAB complex, PolγA and PolγB were mixed in molar ratio of 1:2.

##### Human mtSSB

mtSSB was cloned in pET11a plasmid was expressed in *E. coli* BL21 (DE3) as reported previously (43). Expressed mtSSB was purified by closely following the protocol described by Longley et al. (44) Briefly, mtSSB expressing bacterial cells were lysed using sonication and cell lysate was centrifuged to obtain supernatant with soluble recombinant protein. mtSSB was purified to homogeneity on Blue Sepharose (HiTrap Blue HP), cation exchange (HiTrap SP HP) and anion exchange (HiTrap Q HP) chromatographic columns (all from Cytiva, MA).

##### Twinkle CTD

SUMO-CTD (amino acids 360-684) cloned in pet28 SUMO vector was expressed in *E*.*coli* Rosetta 2 DE3 pLacI cells (50 mg/L Kanamycin). A single colony was grown at 37 °C in 150 mL of LB containing 50 mg/L Kanamycin for 15 h, and inoculated into 1 liter LB containing 50 mg/L Kanamycin, grown at 37 °C to an Optical Density of 0.7, and induced with IPTG (0.2 mM) for ~15 h at 16 °C. The cells were suspended in lysis buffer (50 mM NaH_2_PO_4_ pH 8.0, 1 M NaCl, 1 mM EDTA, 1 mM TCEP, 0.1 % Triton X-100, 0.5 % NP-40, 0.2 mg/mL Lysozyme and Roche protease inhibitor tablets) and sonicated 3 x (amplitude 30, 3 min cycle with 10 s on, 10 s off bursts) on ice, centrifuged (15,500 x g for 1 h) to obtain the supernatant, which was treated with PEI to a final concentration of 0.5 % to remove nucleic acids in the pellet after centrifugation. The proteins in the supernatant were precipitated by ammonium sulfate at 65 % saturation, and resuspended in Buffer A (50 mM NaH_2_PO_4_ pH 8.0, 10 % Glycerol, 0.4 M NaCl, 10 mM Imidazole, 0.1 % Triton X-100, 0.5 % NP-40, 5 mM DTT). The solution was incubated with Qiagen Ni-NTA Agarose bead resin for 1 h, beads were washed 2x with the Buffer A, and spun at 1000 x g for 1 min at 4 °C. The supernatant was removed and protein was eluted with Buffer A containing 200 mM NaCl, 500 mM Imidazole. The eluted protein was mixed with ULP1 SUMO-protease and dialyzed for 12 h in Buffer A with 200 mM NaCl, and applied to a prepacked 1 mL Ni Sepharose High Performance (HP) affinity resin column in the same buffer. The flow-through was applied to a prepacked 1 mL HiTrap Heparin High-Performance column in buffer 50 mM Tris-Cl pH 7.9 10 % Glycerol 200 mM NaCl, 1 mM TCEP and eluted with a 30 ml gradient of 200-1500 mM NaCl. Fractions with pure CTD protein were pooled and concentrated using an Amicon Ultra-2 Centrifugal Filter Unit to ~39 µM of monomeric Twinkle and measured for 260/280 ratio using a Nanodrop (**Fig. S2B Lane 2, D**).

##### Twinkle NTD

SUMO NTD (amino acids 43-372) was cloned into pET 28 expression vector and expressed in BL21 DE3 RIL *E*.*coli* cells using methods described above for the CTD. The lysed cell supernatant was applied to a prepacked 5 mL Ni Sepharose High Performance (HP) affinity resin column in Buffer B (25 mM Tris Cl pH 8.0, 10 % Glycerol, 10 mM Imidazole) containing 400 mM NaCl and eluted using a gradient 250 mL gradient from 0-400 mM Imidazole. The NTD containing fractions were combined and mixed with ULP1 SUMO-protease and dialyzed for 12 h in Buffer B with 200 mM NaCl. The dialyzed solution was applied to a prepacked 1 mL Ni Sepharose High Performance (HP) affinity resin column in Buffer B with 200 mM NaCl, and the flow-through containing cleaved NTD was slowly diluted while stirring at 4 °C with buffer containing 20 mM NaH_2_PO_4_ pH 7.7, 10 mM NaCl, 0.5 mM EDTA, 10 % Glycerol, 1 mM DTT (Buffer C) to bring the NaCl concentration to ~ 100 mM NaCl. The clarified solution applied to a prepacked 1 mL HiTrap Heparin High Performance column in Buffer C with 100 mM NaCl and eluted with 200 ml gradient from 100-415 mM NaCl. The pure NTD fractions were combined and measured to be ~ 26 µM (monomer NTD concentration). (**Fig. S2B Lane 3, E**)

##### FL Twinkle

Full Length Twinkle (aa 43-684) construct was previously prepared in our lab (12). FL Twinkle was expressed as a SUMO fusion in *E*.*coli* Rosetta 2 DE3 pLacI cells, which were grown as described for the CTD, and lysed in Buffer D (50 mM Tris Cl pH 7.1, 600 mM KCl, 1 mM EDTA, 10 % Glycerol, 0.1 % Tween, 0.2 mM TCEP) containing Roche protease inhibitor tablets using a cell homogenizer. The clarified lysate was treated with PEI, and ammonium sulphate as described for CTD, and the ammonium sulphate pellet was dialyzed against Buffer D for 12 h before applying to a prepacked 5 mL Nickel Sepharose High Performance (HP) affinity resin column. Twinkle was eluted using a 40 ml gradient of imidazole from 0-500 mM in Buffer D. The fractions containing SUMO-Twinkle were combined, mixed with ULP1 SUMO-protease, and dialyzed for 12 h in Buffer D with 300 mM KCl, 5 mM EDTA and 5 mM DTT instead of the TCEP and for another 2 h with additional fresh buffer. The dialyzed fractions were applied to a prepacked 1 mL Q Sepharose Fast Flow column in Buffer D containing 150 mM KCl, 5 mM EDTA and 1 mM TCEP and eluted using a 20 mL gradient from 250-1.25 mM KCl. The fractions containing FL Twinkle were applied to a prepacked 1 mL HiTrap Heparin High Performance column in Buffer D with 150 mM NaCl, 5 mM EDTA and 1 mM TCEP and eluted with a NaCl gradient from 150 mM to 1 M. The fractions containing FL Twinkle were measured for 260/280 ratio and those below 0.8 were combined and concentrated using an Amicon Ultra-2 Centrifugal Filter Unit to ~39 µM (monomeric Twinkle concentration). (**Fig. S2B Lane 1, C**)

#### Evaluating Oligomeric State by Gel Filtration Chromatography

Size exclusion chromatography was carried out using Superdex 200 Increase 10/300 GL Cytiva column at 4 °C. Biorad protein standards were applied at 0.3 mL/min in running buffer as described to the chromatography column. The peak elution volumes (280 nm detection) were plotted against the logarithm of protein molecular weight to obtain the calibration curve. Oligomerization of FL Twinkle (15 µl of 38.5 µM monomeric concentration) was tested in Buffer E (50 mM Tris Cl pH 7.1, 5 % Glycerol) with 300 mM NaCl and 1 mM TCEP. Oligomerization of CTD (50 µl of 10 µM monomeric concentration) was tested in Buffer E containing 600 mM NaCl. Oligomerization of NTD (500 µl, 61 µM monomeric concentration NTD) was tested in Buffer C with 200 mM NaCl, and 5 % Glycerol.

#### DNA Binding Using Fluorescence Anisotropy Titrations

Serial diluted protein solution was mixed with fluorescein-labeled DNA (IDT, HPLC purified) at 2.5 nM concentration in buffer containing 50 mM Tris acetate pH 7.5, 10 % glycerol, 0.05 % Tween 20, 0.5 mM DTT with and without 50 mM NaCl. Fluorescence intensities were measured on the TECAN Spark plate reader at excitation wavelength 485 (20 nm bandwidth) and emission wavelength 535 (20 nm bandwidth). The anisotropy (*r*) was calculated from the parallel and perpendicular polarized light emission intensity (*I*) utilizing the equation: 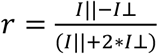 and plotted against protein concentration (*P*)(monomer for NTD, hexamer for CTD and FL Twinkle). The curves were fit to equation as follows: 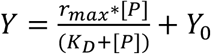 or each binding titration curve with the exception of the NTD binding to 30 nt and 20 nt length ssDNAs in the absence of salt which were fit the equation as follows: 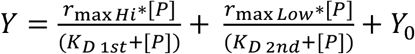 Here, *Y*_*0*_ is the anisotropy of free DNA, *rmax* is the anisotropy of protein-bound DNA.

#### Radiometic Assay to Measure ATP hydrolysis

The ATPase reactions were carried out in Buffer F (50 mM Tris acetate pH 7.5, 0.01 % Tween 20, 1 mM EDTA, 5 mM DTT) with and without 2.5 nM M13 ssDNA using 30 nM Twinkle hexamer, 8 mM Mg acetate, and 1 mM ATP spiked with [γ^32^P]ATP. Reactions were quenched with 8 M formic acid, spotted on a PEI cellulose TLC, and developed in 0.4 M potassium phosphate pH 3.4. The counts from the resolved ATP and inorganic phosphate spots were used to obtain the proportion of ATP hydrolysis with time.

#### DNA Unwinding Assay

DNA unwinding assays were carried out in Buffer F using 10 nM preannealed fluorescein-labeled DNA fork, 55.5 nM FL Twinkle or CTD hexamer, 4.5 mM ATP and 8 mM magnesium acetate. Buffer components, ATP, DNA fork, and Twinkle are combined in solution A, and magnesium acetate and trap DNA (unlabeled upper strand, 280 nM final concentration) were combined in solution B. solution A and B were mixed in equal volumes to begin the reaction at 30 °C. Portions of the reaction were removed at various time intervals, diluted 1:1 with quenching solution containing 100 mM EDTA pH 8.0 and 1 % SDS to stop active enzymatic activity and denature the protein. The quenched samples were mixed 1:10 with loading buffer (15 % ficoll 400 in 0.3x TBE) loaded into a 4-20 % TGX gradient gel running at a low voltage of 150 V while loading samples. The gel was run for approximately 2 h at 4 °C until the bromophenol blue dye in an otherwise empty lane reached near the bottom of the gel but did not run off. The gels were scanned for fluorescence on a GE Typhoon FLA 9000 Gel Scanner at 600 V in the FAM mode and processed using ImageQuant TL 8.2 in 1D gel analysis mode. The rolling ball method of background subtraction was used on the least sensitive (200) setting, background subtracted counts were graphed and analyzed in GraphPad Prism 9.0.

#### Strand Displacement DNA Synthesis on Short Replication Fork

Reactions contained 100 nM assembled primed fork DNA, 200 nM wild-type Polγ (A and B in 1:2 ratio), 300 nM FL Twinkle or CTD hexamer, 10 mM MgCl_2_, 4 mM ATP, 100 µM dNTPs, 300 µM dCTP in buffer containing 50 mM Tris Cl pH 7.5, 40 mM NaCl, 10 % glycerol, and 2 mM DTT. Buffer components, Twinkle, Polγ, fork DNA, dCTP were combined into solution A over a series of additions and incubations. Polγ was added to primed fork DNA with dCTP (next nucleotide) and incubated on ice for 5 min in buffer without MgCl_2_. Twinkle or CTD was added, incubated on ice for 20 min, and then at 37 °C for 5 min before adding dNTPs, MgCl_2_ and ATP. Portions of reaction were removed and quenched at 5, 15, 30 and 60 min with 500 mM EDTA, 2 μM trap DNA (unlabeled upper strand). Samples were diluted 1:1 with 100 % formamide, boiled for 5 min before being quickly transferred to ice and loaded into a 15 % TBE-Urea denaturing gel and run at 250 V until bromophenol blue loaded into an otherwise empty lane migrated 2/3 of the way through the gel. Polγ incubated with the primer annealed only to the complementary lower strand without the upper strand present was used as a positive control for full extension, while the primer alone was used for a no extension control. The gels were scanned and analyzed as above.

#### Rolling Circle DNA Synthesis Assay

Reaction contained 20 nM Polγ, 10 nM 70 bp Minicircle fork, 250 nM mtSSB tetramer, 20 nM FL Twinkle or CTD hexamer, 10 mM MgCl_2_, 2 mM ATP, 0.25 mM dATP, 0.25 mM dCTP, 0.25 mM dTTP, and 25 µM dGTP spiked with [α-^32^P]dGTP in 50 mM Tris Cl pH 7.5, 10 % Glycerol, 2 mM DTT. Buffer components, minicircle fork, Polγ, and mtSSB were combined, in that order in solution A and allowed to incubate for 20 min at 37 °C. MgCl_2_, ATP, dNTPs and [α-^32^P]dGTP were combined in solution B and added to solution A to begin the reaction. Portions of the reaction were removed at various time intervals, diluted 1:1 with quenching solution containing 100 mM EDTA pH 8.0 to stop active enzymatic activity. Proteinase K was added to a final concentration of 0.1 mg/mL and the samples incubated for 45 min at 42 °C to digest any bound protein off of the DNA. The digested samples were mixed 1:10 with loading buffer (300 mM NaOH, 6 mM EDTA, 18 % (w/v) ficoll 400, 0.15 % (w/v) bromophenol blue and 0.25 % xylene cyanol). The samples were loaded on 0.8 % alkaline agarose gel with 1 kb NEB ladder labeled with [γ-^32^P] ATP using T4 polynucleotide kinase, and electrophoresed in running buffer containing 50 mM NaOH, 6 mM EDTA at 74 V and 4 °C for 15.5 h. The gel was fixed in 7 % TCA, dried and exposed to a phosphorscreen, scanned on a Typhoon scanner, and analyzed using ImageQuant software.

## Data Availability

All the data are in the manuscript.

## Acknowledgements

We would like to thank the Patel Lab members for advice and suggestions on this work.

## Author contributions

L.J., A.S., and S.S.P. conceptualization, data curation, formal analysis, and S.S.P. funding acquisition; project administration; L.J. original draft, and S.S.P. writing and editing.

## FOOTNOTES

Funding for this work was provided by the National Institute of General Medical Sciences (NIGMS) [GM118086] to S.S.P.

## SUPPORTING INFORMATION (FIGURES AND TABLES)

**Figure S1.**
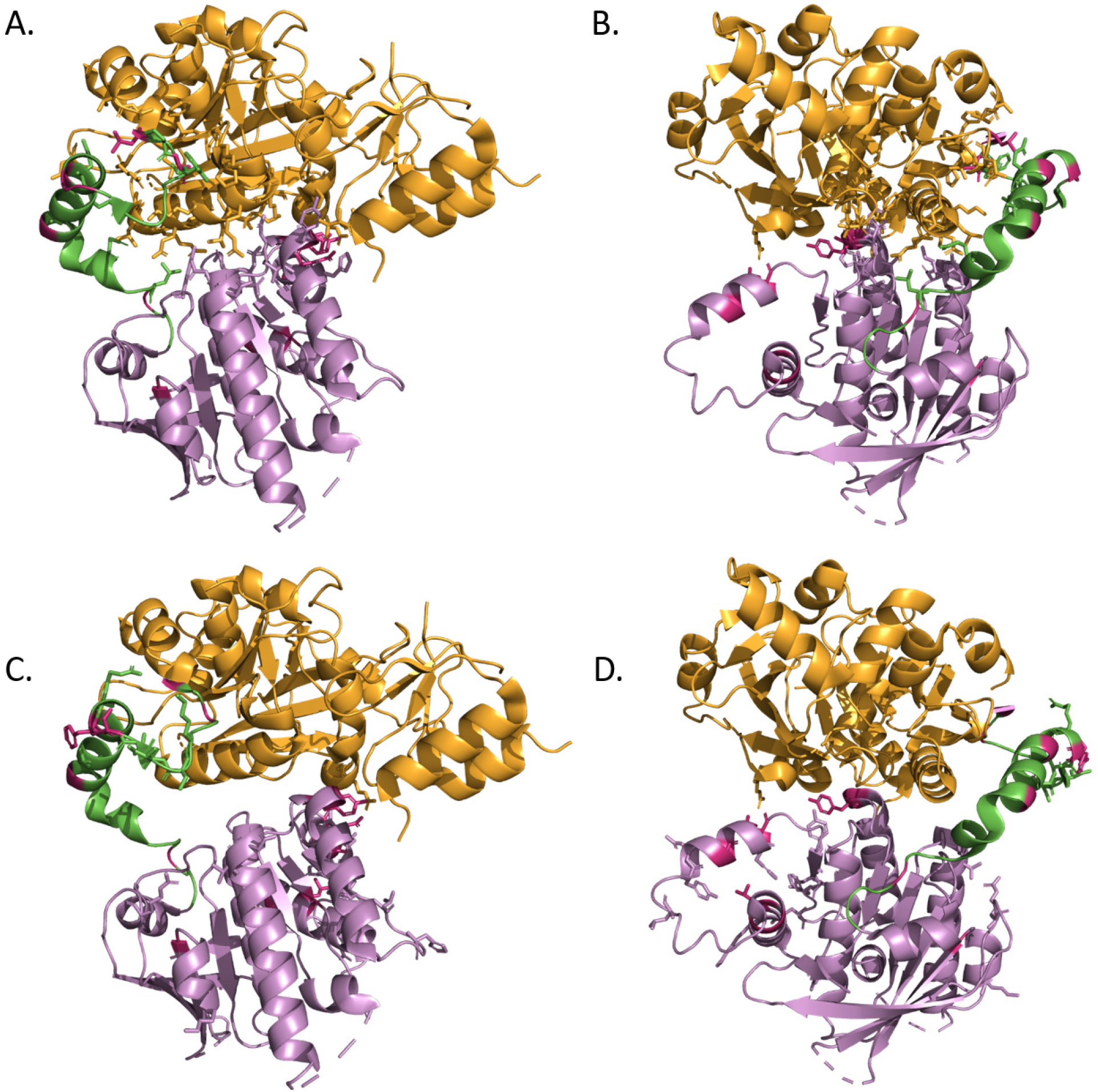
Isolated subunit of the heptameric Twinkle (PDB ID: 7T8C) with residues (sticks) involved in inter- and intra-molecular interactions in Twinkle highlighted within 4 Angstroms and locations of the human clinical disease mutants (in fuchsia). The NTD is displayed in orange, the CTD in purple and the linker in green. **Panels A and B** show different orientations of one subunit of FL Twinkle with NTD-CTD intramolecular interface residues displayed as sticks. The NTD interacts extensively with the linker region involved in subunit interactions. **Panels C and D** show different orientations of one subunit of FL Twinkle with intermolecular interface residues displayed as sticks. Most of the intermolecular interactions between subunits lie in the linker and CTD.

**Figure S2.**
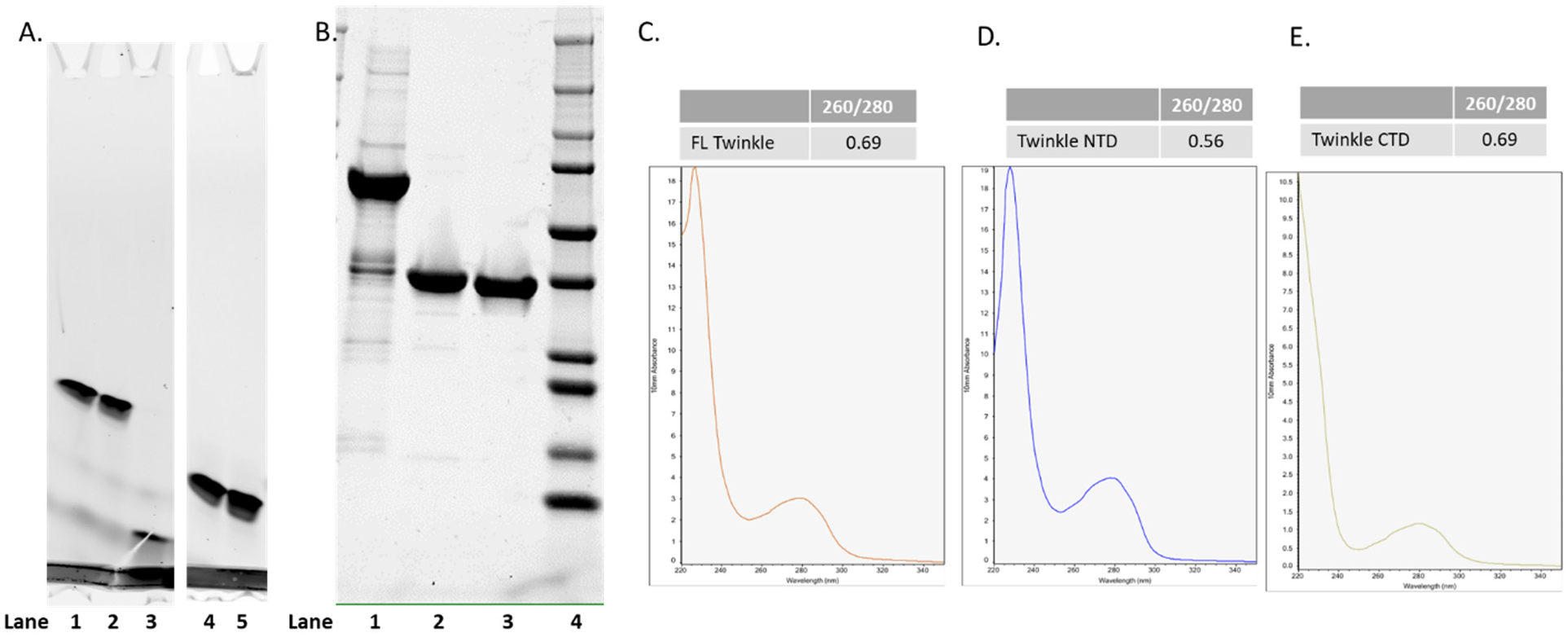
Purified twinkle constructs quality assessment **A**. Gel analysis of nuclease contamination in different Twinkle preparations. 5 nM 35-mer ssDNA labeled with 6-FAM at the 3’ end was incubated with 300 nM (hexamer) of insect cell expressed Twinkle purified on Heparin column or Twinkle expressed in *E. coli* and purified with or without Heparin column step. 10 mM MgCl_2_ was added to the DNA or DNA-Twinkle complexes to initiate the reactions. Reactions were stopped after 30 minutes with 100 mM EDTA and 0.5 % SDS and equal volumes were loaded and analyzed on 4-20 % SDS-PAGE gel. Gel was scanned on FLA Typhoon 9500 scanner (GE Healthcare) to visualize the fluorescent DNA bands. Intact DNA in reactions conducted with Twinkle preparations including Heparin column step confirms that the step removed the contaminating nuclease(s). All the reactions were conducted at 25°C. Lane 1: DNA, Lane 2: DNA + Insect cell expressed FL Twinkle (after Heparin column), Lane 3: DNA + *E*.*coli* cell expressed FL Twinkle FL (no Heparin column) with strong visible exo activity, Lane 4: DNA, Lane 5: DNA + *E*.*coli* cell expressed FL Twinkle (after Heparin column). **B**. 4-20 % SDS-PAGE gel of all three purified Twinkle constructs appearing < 95 % pure of other protein contaminants, Lane 1: FL Twinkle, Lane 2: Twinkle CTD, Lane 3: Twinkle NTD, Lane 4: BioRad Precision Plus Protein™ Ladder **C**. Twinkle absorbance spectra and 260/280 ratio **D**. Twinkle CTD absorbance spectra and 260/280 ratio **E**. Twinkle NTD absorbance spectra and 260/280 ratio

**Figure S3.**
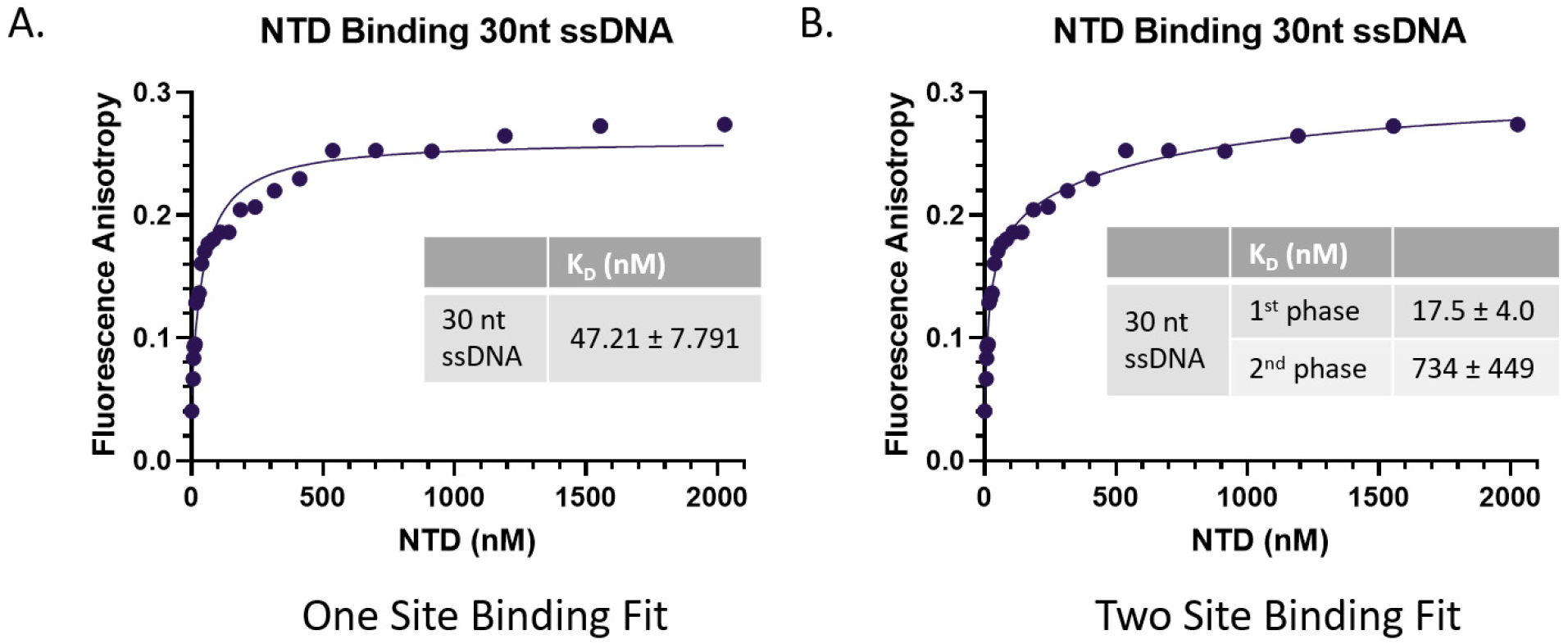
Comparison of binding titration curves of Twinkle NTD binding 30 nt ssDNA fit to either one or two site hyperbolic binding equation, the curve exhibiting clear two phase character suggesting a tight initial binding and a subsequent weaker binding of an additional NTD subunit **A**. one site binding hyperbolic equation Y = r_max_*[P]/(K_D_ + [P])+ Y_0_ **B**. two site binding hyperbolic equation Y = r_maxHi_*[P]/(K_D 1st_ + [P]) + r_maxLow_*[P]/(K_D 2nd_ + [P]) + Y_0_. Here, r_max_ is the maximum anisotropy, [P] is protein concentration, Y_0_ is y-intercept.

**Table S1.**
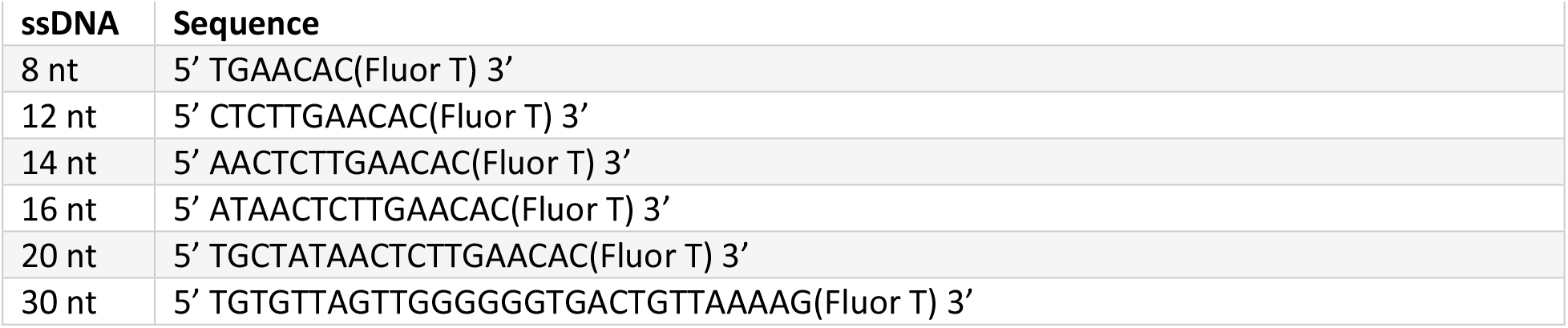
ssDNA length series constructs used in fluorescence anisotropy binding experiments

**Table S2.**
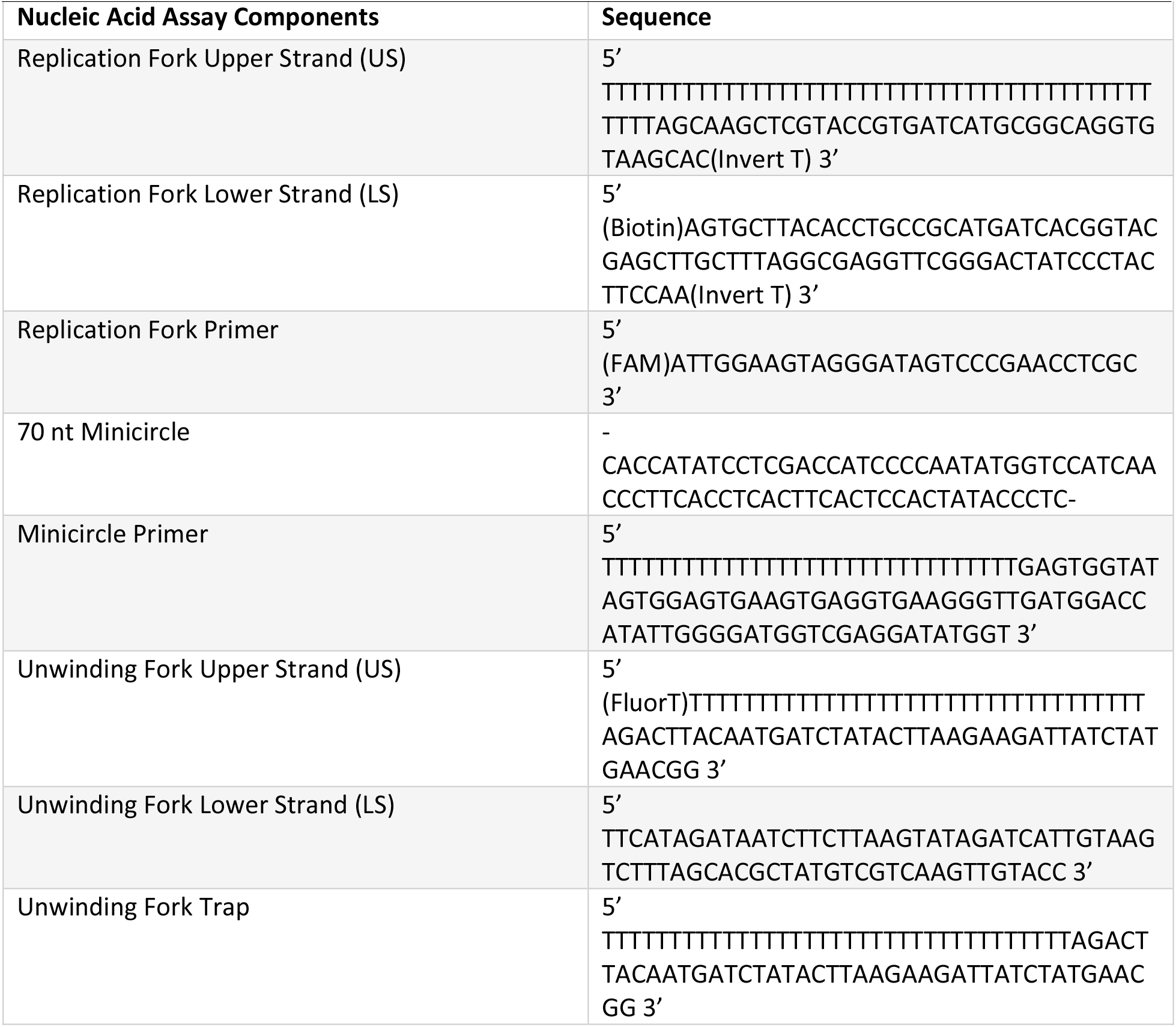
DNA construct components used in short fork strand displacement synthesis, helicase unwinding, and rolling circle synthesis experiments

**Table S3.** Total rolling circle product counts quantified from area under the curve for the product counts over length graphs

|  |  | **Product**<br>**(Counts)** |
| --- | --- | --- |
| FL | 60 min | 148013 |
|  | 60 min + mtSSBs | 334720 |
| | 60 min + ATPγS | 7285 |
| | 60 min + ATPγS + mtSSBs | 33632 |
| CTD | 60 min | 5468 |
|  | 60 min + mtSSBs | 68547 |
| | 60 min + ATPγS | 5448 |
| | 60 min + ATPγS + mtSSBs | 26701 |

